# Host-Microbial Interactions in Systemic Lupus Erythematosus and Periodontitis

**DOI:** 10.1101/631051

**Authors:** L.C. Pessoa, G. Aleti, S. Choudhury, D. Nguyen, T. Yaskell, Y. Zang, L. Weizhong, K.E. Nelson, L. L. Santos Neto, A.C.P. Sant’Ana, M. Freire

**Affiliations:** Department of Genomic Medicine, J. Craig Venter Institute, 4120 Capricorn Lane, La Jolla, CA 92037, USA; Department of Genomic Medicine, J. Craig Venter Institute, 9605 Medical Center Drive Suite 150, Rockville, MD 20850, USA; Department of Prosthodontics and Periodontics, Bauru School of Dentistry, University of São Paulo, Bauru, SP, Brazil. Al. Dr. Octávio Pinheiro Brisolla 9-75, 17012-101. Bauru, SP, Brazil; Applied Oral Sciences, The Forsyth Institute, 245 First Street, Cambridge, MA 02142, USA; Rheumatology Department, Medical Faculty, University of Brasília. 70910-900, DF, Brazil

**Keywords:** Cytokines, Gingival crevicular fluid, Subgingival microbiota, Periodontitis

## Abstract

Systemic lupus erythematosus (SLE) is a potentially fatal complex autoimmune disease, that is characterized by widespread inflammation manifesting tissue damage and comorbidities across the human body including heart, blood vessels, joints, skin, liver, kidneys, and periodontal tissues. The etiology of SLE is partially attributed to a deregulated inflammatory response to microbial dysbiosis and environmental changes. In the mouth, periodontal environment provides an optimal niche to assay local dynamic microbial ecological changes in health and disease important to systemic inflammation in SLE subjects. Our aim was to evaluate the reciprocal impact of periodontal subgingival microbiota on SLE systemic inflammation. Ninety-one female subjects were recruited, including healthy (n=31), SLE-inactive (n=29), and SLE-active (n=31). Patients were screened for probing depth (PD), bleeding on probing (BOP), clinical attachment level (CAL), and classified with or without periodontal dysbiosis, periodontitis. Serum inflammatory cytokines were measured by human cytokine panel and subgingival biofilm was examined by DNA-DNA checkerboard. The results showed significant upregulation of proinflammatory cytokines in individuals with SLE when compared to controls. Stratification of subject’s into SLE-inactive (I) and SLE-active (A) phenotypes or periodontitis and non-periodontitis groups provided new insights into SLE pathophysiology. While low-grade inflammation was found in SLE-I subjects, a potent anti-inflammatory cytokine, IL-10 was found to control clinical phenotypes. Out of twenty-four significant differential oral microbial abundances found in SLE, fourteen unique subgingival bacteria profiles were found to be elevated in SLE. Pathogens from periodontal disease sites (*Treponema denticola* and *Tannerella forsythia*) showed increase abundance in SLE-A subjects when compared to controls. Cytokine-microbial correlations showed that periodontal pathogens dominating the environment increased proinflammatory cytokines systemically. Deeper clinical attachment loss and periodontal pathogens were found in SLE subjects, especially on SLE-I, likely due to long-term chronic and low-grade inflammation. Altogether, local periodontal pathogen enrichment was positively associated with high systemic inflammatory profiles, relevant to the overall health and SLE disease pathogenesis.

## 1. Introduction

The human microbiome is in constant interaction with the host, modulating health and disease phenotypes. We now appreciate that oral bacteria play indispensable roles in shaping the systemic host physiological landscape and in dysbiosis. As part of the upper digestive tract, the oral cavity presents specific niches, such as gingival sulcus and gingival crevicular fluid (GCF), which in turn harbor commensal and pathogenic bacteria with potential impact to oral and systemic health. Through local activation of inflammation, oral pathogens have shown to worsen the burden of chronic diseases through time including, type 2 diabetes, premature labor, rheumatoid arthritis, systemic lupus erythematosus (SLE), Alzheimer’s, cardiovascular conditions, and cancer ^1–4^.

Locally, the oral commensal flora and tissue inflammation evolved to develop a relationship of homeostasis. In dysbiosis, pathogens become dominant, including species previously clustered according to their pathogenesis into the “red complex”-*Tannerella forsythia*, *Porphyromonas gingivalis*, and *Treponema denticola ^5^.* In periodontitis, for example, continuous and unresolved inflammatory response affects tooth-supporting tissue structures (periodontal ligament, connective tissue, and bone) ^3,6^. As one of the most common infectious diseases globally, the etiology of periodontitis is multifactorial, and the microbial-inflammation imbalance leads to bone resorption and consequent tissue tooth loss ^7^. Inflammation precedes tissue loss and the systemic impact of periodontal dysbiosis goes beyond the oral compartment.

Systemic lupus erythematosus (SLE) is a multisystem autoimmune disease with increasing incidence worldwide ^8^. The Lupus foundation of America estimates that 1.5 million Americans have SLE, and at least five million people worldwide have a form of lupus ^9,10^. SLE affects mostly women and most people with the condition manifests the symptoms between ages of 15-44. The current incidence of SLE is 16,000 new cases per year and patients experience significant symptoms, such as pain, and fatigue, hair loss, cognitive issues, physical impairments, oral and vaginal mucosa manifestations, affecting every facet of their lives ^11^. According to the American College of Rheumatology (ACR), an array of clinical exams and positive serology are used to diagnose disease, including malar rash, discoid rash, arthritis, kidney disorder, anemia or leukopenia, abnormal antinuclear antibodies (ANA) and anti-DNA or antiphospholipid antibodies ^12^. In addition to molecular diagnosis, systemic lupus erythematosus disease activity index (SLEDAI) is used in the assessment of disease severity and response to treatment; manifestations of comorbidities and molecular abnormalities affecting multiple organs of the body often challenges diagnostics at early stages ^13^. Although complex diseases have shown to be influenced by genetic and environmental forces, the gut microbiome, and recently the oral microbiome, showed direct impact on SLE subjects ^14^, the complexity of chronic diseases such as SLE is beyond isolated body compartments, and it requires the integration of host-microbial interactions. The heterogeneity of disease presentation and organ involvement contribute to clinical challenges for diagnosis and effective management ^15^. While several studies reported associations among human oral microbiota compositions in SLE ^16–19^, co-occurrences of specific periodontal pathogens and inflammatory cytokines important for low grade inflammation and chronic disease remains to be explored ^4^.

In this study we have investigated SLE phenotypic differences, oral microbiota associations with systemic inflammation to unravel forces shaping oral and broader systemic health. As such, we compared the subgingival microbial signatures from SLE and healthy controls to investigate host-microbial relationships. Specifically, we first sought to establish if specific microbial compositions correlated to periodontal clinical phenotypes with systemic levels of inflammatory cytokines. The presence of periodontal disease was assessed using categories of local inflammation, clinical attachment levels and bone loss, while SLEDAI > or < 2 was assessed to categorize the individuals into SLE-active, SEL-inactive. The results indicate that chronic periodontitis and SLE present low-grade inflammation modulated periodontal diseases and specific microbial signatures. The association of subgingival microbial profiles with SLE and its association with periodontal clinical status and inflammatory markers established novel links developing a new framework for oral-systemic studies.

## 2. Materials and Methods

### 2.1 Subjects

Subjects were recruited at the Center of Rheumatology at the University Hospital in Brasília from July 2013 to March 2014, and samples were obtained with informed consent of the subjects. The diagnosis of SLE was initially made by the primary care physician and confirmed by rheumatologist specialist following guidelines of the American Academy of Rheumatology (according to ACR 1982/1997 revised classification criteria) ^20,21^. This research project was approved by the Bauru Dental School-USP Ethics Committee of Research (protocol # 111.718) and by the Ethics Committee of Research. Subjects with antimicrobial treatments within 3 months of the start of the study were excluded. Clinical parameters for SLE and periodontal status were measured as follows:

#### SLE status

SLE activity and diagnosis was investigated according to SLEDAI classification ^22^. SLE activity was determined by SLEDAI > 2. Disease severity is measured according to SLICC/ACR-DI (Systemic Lupus International Collaborating Clinics of American College of Rheumatology Damage Index) proposed by Gladman et al. ^23^, which was used as an additional method of information on disease severity to better characterize the studied population. SLE patients were subdivided into two groups based on their disease activity: active (SLE-A; SLEDAI > 2; n= 29), and inactive (SLE-I; SLEDAI ≤ 2; n=31). The control group was composed of 31 systemically healthy women recruited at the University Hospital in Brasília during the same time period. All patients answered a health questionnaire investigating medical and dental history. A visual investigation of the oral cavity was performed at enrollment to investigate the number of remaining teeth, other oral diseases and periodontal health.

#### Periodontal Status

After recruitment of subjects, participants were referred to the Dental School of Brasília (Brasilia, Brazil) for clinical periodontal examination and sample collection. The clinical periodontal examination was performed by a single trained examiner before the collection of GCF, subgingival plaque and blood samples prior to oral and non-surgical periodontal treatment. The following parameters were investigated by using a UNC-15 millimeter periodontal probe at 6 sites/tooth: pocket probing depth (PD), clinical attachment level (CAL) and gingival bleeding index ^24^. Plaque index was assayed by staining the dental plaque with disclosing solution at mesial, buccal, distal and lingual sites ^25^. After periodontal examination, periodontal disease was classified according to CDC and AAP criteria ^25–27^ including: A) Mild periodontitis: 2 or more interproximal sites with ≥ 3 mm of CALs and ≥ 4 mm PD (not on the same tooth) or 1 site with PD ≥ 5mm; B) Moderate periodontitis: 2 or more interproximal sites with ≥ 4 mm of CAL (not on the same tooth) and 2 or more interproximal sites ≥ 5 mm PD (not on the same tooth); C) Severe periodontitis: 2 or more interproximal sites with ≥ 6 mm of CALs (not on the same tooth) and 1 or more interproximal site(s) with ≥ 5 mm PD. Total prevalence of periodontitis was determined by the sum of mild, moderate and severe periodontitis. The extent of disease was reported by the severity of disease at 5, 10 and 30% of sites and teeth. After radiographic and clinical examination followed by collection of blood, GCF and subgingival plaque samples, patients were submitted to non-surgical periodontal treatment consisted of supra and subgingival scaling and root planing, when necessary, dental polishing and oral hygiene instruction.

### 2.2 Sample Collection

Subgingival plaque and GCF sample collection were collected prior to any treatment for all subjects, including non-surgical periodontal treatment. GCF and subgingival plaque samples were collected from four sites showing the deepest probing depth in each patient. The area was isolated with cotton rolls to prevent contact with saliva and air dried. Supragingival plaque was removed with Gracey curettes previous to subgingival plaque collection. The subgingival plaque was retrieved by the introduction of PerioPaper strips into gingival sulcus for 30 seconds and stored in sterilized tubes at −80°C till processing. Blood samples were collected after routine examination for SLE monitoring. Control subjects were asked to perform similar examinations at the Hospital. The samples were stored at −80°C up to processing. PerioPaper strips were centrifuged at 3000 × g for 15 min at 4°C in PBS elution for the dosage of the tissue destruction and bone resorption markers by Multiplex (Human MAP, Millipore, USA).

### 2.3 DNA extraction, Amplification and Checkerboard measurements

DNA-DNA checkerboard hybridization was performed using the method developed by Socransky et al. (1998). Briefly, to prepare standard probes, forty selected bacterial species were grown on agar plates and were suspended in 1 mL of TE buffer (10 mM Tris-HCl, 0.1 mM EDTA, pH 7.6). Cells were pelleted at 1300 × g for 10 min and washed with TE buffer. Gram-negative strains were lysed with 10% SDS and Proteinase K (20 mg/mL) while Gram-positive strains were lysed with 15 mg/mL lysozyme (Sigma) and 5 mg/mL achromopeptidase (Sigma). Following, cells were sonicated and incubated at 37°C for 1 hour. DNA was isolated and purified as described previously by Smith et al. (1989). DNA probes were generated for each strain by labeling with digoxigenin (Roche, Basel, Switzerland) ^28^. Next, the DNA content of the subgingival plaque samples was amplified using multiple displacement amplification ^29^. Briefly, subgingival DNA templates were mixed with random hexamer primers in 50 mM Tris-HCl (pH 8.2) and 0.5 mM EDTA. The solution was heat denatured at 95°C for 3 min in a thermocycler. DNA polymerase with dNTPs was then added and incubated at 30°C for 2 hours. The subgingival plaque samples along with 1 ng and 10 ng of known DNA standards were analyzed by using checkerboard DNA-DNA hybridization as previously described by Socransky *et al*., (2004). In brief, samples were boiled for 10 min in 1 mL of TE buffer and were loaded on a nylon membrane (Roche, Basel, Switzerland) using a Minislot 30 apparatus (Immunetics, Cambridge, MA, USA). Then cross-linked by ultraviolet light using UV-crosslinker (Stratalinker 1800, La Jolla, CA, USA). Probes were subsequently bound perpendicular to the samples on the membrane and were detected using anti-digoxigenin antibody conjugated with alkaline phosphatase and a chemifluorescent substrate. Signal intensities of the plaque samples were measured using a Typhoon TRIO+ Scanner (GE Healthcare). Finally, absolute counts were generated by comparing with signal intensities of standards using Phoretix Array Software (TotalLab) ^30,31^.

### 2.4 Principal Coordinate analysis

The differences between control and diseased groups were investigated by applying Principal Coordinates Analysis (PCoA) on the binary Jaccard distance metric using ClusterApp tool (available at http://dorresteinappshub.ucsd.edu:3838/clusterMetaboApp0.9.1/) and visualized via EMPeror software ^32^. Pairwise permutation MANOVA (ADONIS within RVAideMemoire package in R) with 999 permutations were used to test significant differences between sample groups.

### 2.5 Phylogenetic analysis

The 16S rRNA gene sequences were downloaded from the Human Oral Microbiome Database (HOMD; version 14.51, January 2017), and were aligned using ClustalW software within Molecular Evolutionary Genetics Analysis (MEGA) software version 7^33^. The phylogenetic tree was plotted by applying the Neighbor-Joining method using p-distance model and by implementing 1,000 bootstrap replications. Species abundance data and the phylogenetic tree were jointly visualized further using the interactive tree of life (iTOL) version 4 (available at http://itol.embl.de/) ^34^.

### 2.6 Cytokines measurement

Blood plasma collected from the subjects were immediately stored at −80℃. A panel of 20 inflammatory cytokines were measured in duplicate by ProcartaPlex Human Inflammation Panel 20-plex (ThermoFisher Scientific, San Diego, USA) on a Luminex 200 System (Luminex Corporation, Texas, USA) as previously described ^35^ and adapted according to the manufacturer’s specifications. Positive and negative controls were assayed on each plate. The cytokines measured were macrophage inflammatory protein 1 alpha (MIP-1-ɑ), interleukin (IL)-1 beta (β), IL-4, IL-6, IL-8, IL-10, IL-12p70, IL-13, IL-17A, IL-1ɑ, interferon gamma-Induced Protein 10 (IP-10), interferon gamma (IFN-ɣ), granulocyte-macrophage-colony-stimulating factor (GM-CSF), tumor necrosis factor alpha (TNF-ɑ), monocyte/macrophage inflammatory protein-1 beta (MIP-1β), interferon alpha (IFN-ɑ), monocyte chemoattractant protein 1 (MCP-1), Platelet-selectin (P-selectin), soluble intercellular adhesion molecule-1 (sICAM-1), Endothelial-selectin (E-selectin). Cytokine abundance levels were reported in pg/mL by the assay system. Quality control of the machine-generated raw data was performed using xPONENT 4.2 software as per manufacturer’s guidelines (Affymetrix eBioscience, San Diego, USA).

### 2.7 Statistical analyses

Statistical analyses were applied to the microbial and cytokine abundance data to detect any features that are significantly different between diseased and control groups. Rank-based tests were employed as a robust nonparametric group comparison tool for those non-normally distributed microbiome and cytokine data. Using Wilcoxon rank-sum test, we further analyzed differences between groups i.e., control versus SLE-I, control versus SLE-A and SLE-I versus SLE-A. For significance testing, FDR-corrected (Benjamini-Hochberg) p-values were set at 5% threshold. For binary abundances we dichotomized the relative abundances data into presence (i.e. abundance > 0) and absence (i.e. abundance = 0) binary values. We used a logistic regression model to analyze the relationship between the binary abundance data and two clinically important categorical covariates: SLE group and periodontal condition^36^. For the disease group, we have three levels (control, SLE-A, and SLE-I); for the periodontal condition, we have two levels (control versus periodontitis).

Potential associations between microbial species and cytokine abundances were evaluated by Spearman’s rank-based correlations. Spearman’s correlation coefficients were calculated using R package *psych*, and significant correlations were plotted with R package *corrplot*. Pairwise correlations were computed separately for control and SLE samples. To avoid false positives, we selected correlations with adjusted p-values <0.05. For visual simplicity, we showed only the significant correlations. Topological parameters were analyzed by importing the pairwise correlations using network analyzer algorithm within Cytoscape, 3.7.1. Hierarchical clustering was applied on the relative abundance data of species in each sample based on Spearman rank correlation and the heatmap was generated using *ggplots* package in R.

## 3. Results

### 3.1 Clinical Findings of the Cohort

To investigate the role of the periodontal microbiome-induced inflammation on SLE subjects, whole blood, crevicular fluids, and plaque were collected from subjects enrolled in the study. General characteristics of patients are shown in Table 1. A total of 91 subgingival samples were analyzed from women aged 18-65. SLE activity was determined by SLEDAI > 2 ^37^. Of 91 subjects, 31 individuals were controls, 29 were SLE-I and 31 were SLE-A. The average SLE disease activity index (SLEDAI) for SLE-A subjects significantly increased (7.29 ± 4.31), while SLE-I showed a lower SLEDAI (1.07 ± 0.99). Patients with SLE-I had a longer duration of the disease and aged significantly older, whereas the SLE-A patients were relatively younger than controls and SLE-I subjects (32.58 ± 8.9 years, p = 0.045). SLE-I individuals showed increased clinical attachment loss (CAL) and worsen periodontal clinical outcomes (8.31 ± 9.438). The majority of oral health indices including teeth number, bleeding points, BOP (Bleeding on probing), stained teeth, plaque index, and probing depth were similar between SLE groups. Ethnicity was equally distributed among the three groups of the studied population.

**Table 1.**
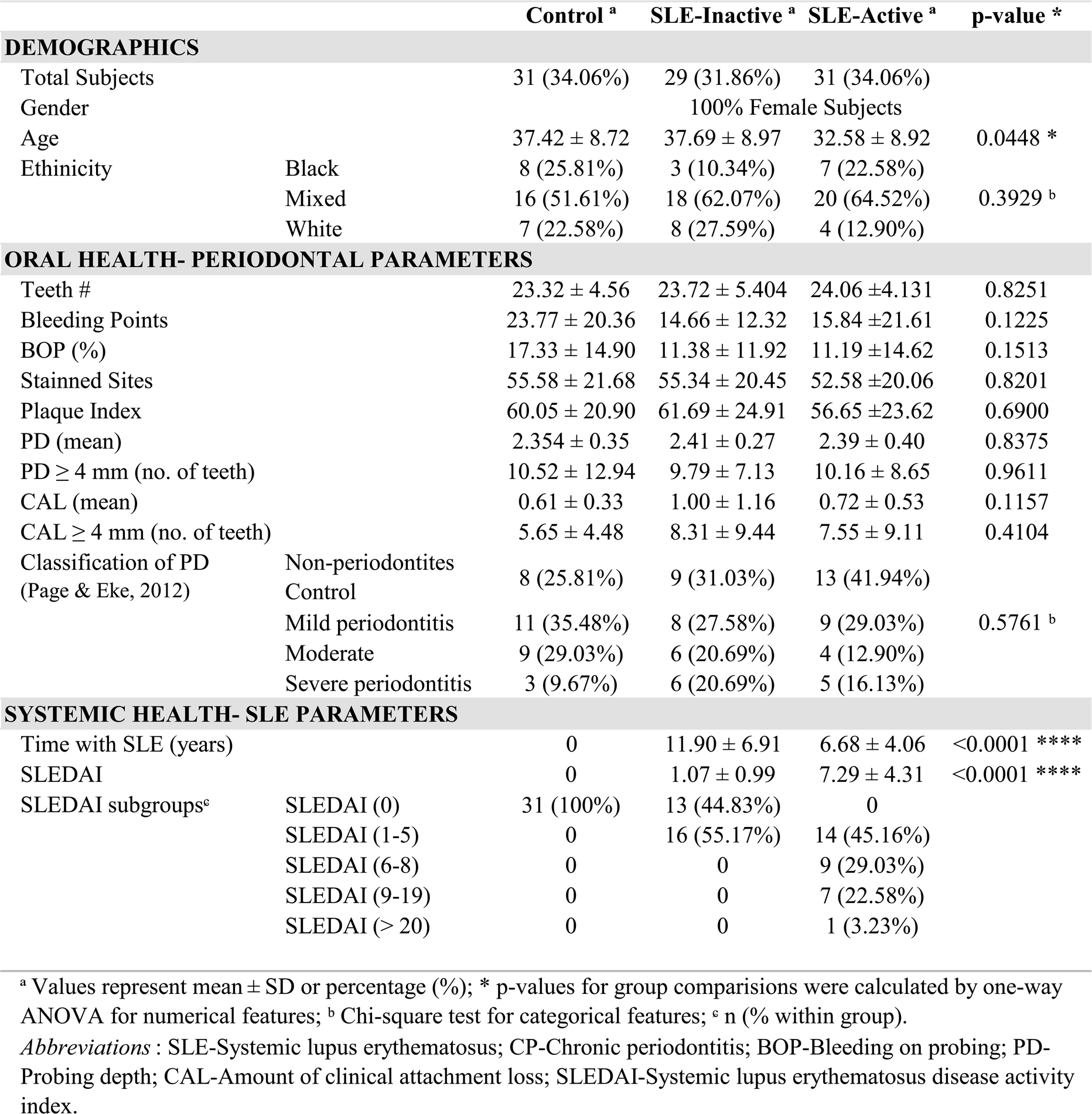
Demographic and clinical features of SLE and control subjects.

### 3.2 Inflammatory Cytokines Upregulation in SLE

Various inflammatory cytokines have been involved in regulating disease activity and in organ pathologies of patients diagnosed with SLE ^38,39^. In order to identify specific markers responsible for systemic inflammation in SLE patients, we investigated whole blood serum concentrations of pro and anti-inflammatory cytokines from SLE groups compared to healthy control. We measured proinflammatory levels of MIP-1-ɑ, IL-1β, IL-4, IP-10, IL-6, IL-8, IL-12p70, IL-17A, IFN-ɣ, GM-CSF, TNF-ɑ, MIP-1β, IFN-ɑ, MCP-1, P-Selectin, IL-1ɑ, sICAM-1, E-selectin, and anti-inflammatory cytokines IL-10, IL-13. Our results demonstrated that twenty cytokine concentration comparisons were upregulated in SLE subjects when compared to healthy controls (Figure 1A, Supplementary Table 1), with sixteen unique profiles among SLE groups (Figure 1B). Eleven proinflammatory cytokine profiles were upregulated in SLE-I only (IFN-ɑ, IL-17A, MIP-1-ɑ, IL-4, IFN-ɣ, IL-10, siCAM-1, GM-CSF, IL-12p70, IL1β, IL-1ɑ), and E-selectin was unique to SLE-A only (Figure 1B). SLE-I sera also demonstrated anti-inflammatory increase by upregulation of IL-10, this was not evident on SLE-A. Among the significant differences from SLE, inactive and active groups, there were four molecules that overlapped (MCP-1, IL-8, IP-10, IL-6). These results suggest that increased low-grade systemic inflammation was observed in SLE patients when compared to healthy controls.

**Figure 1.**
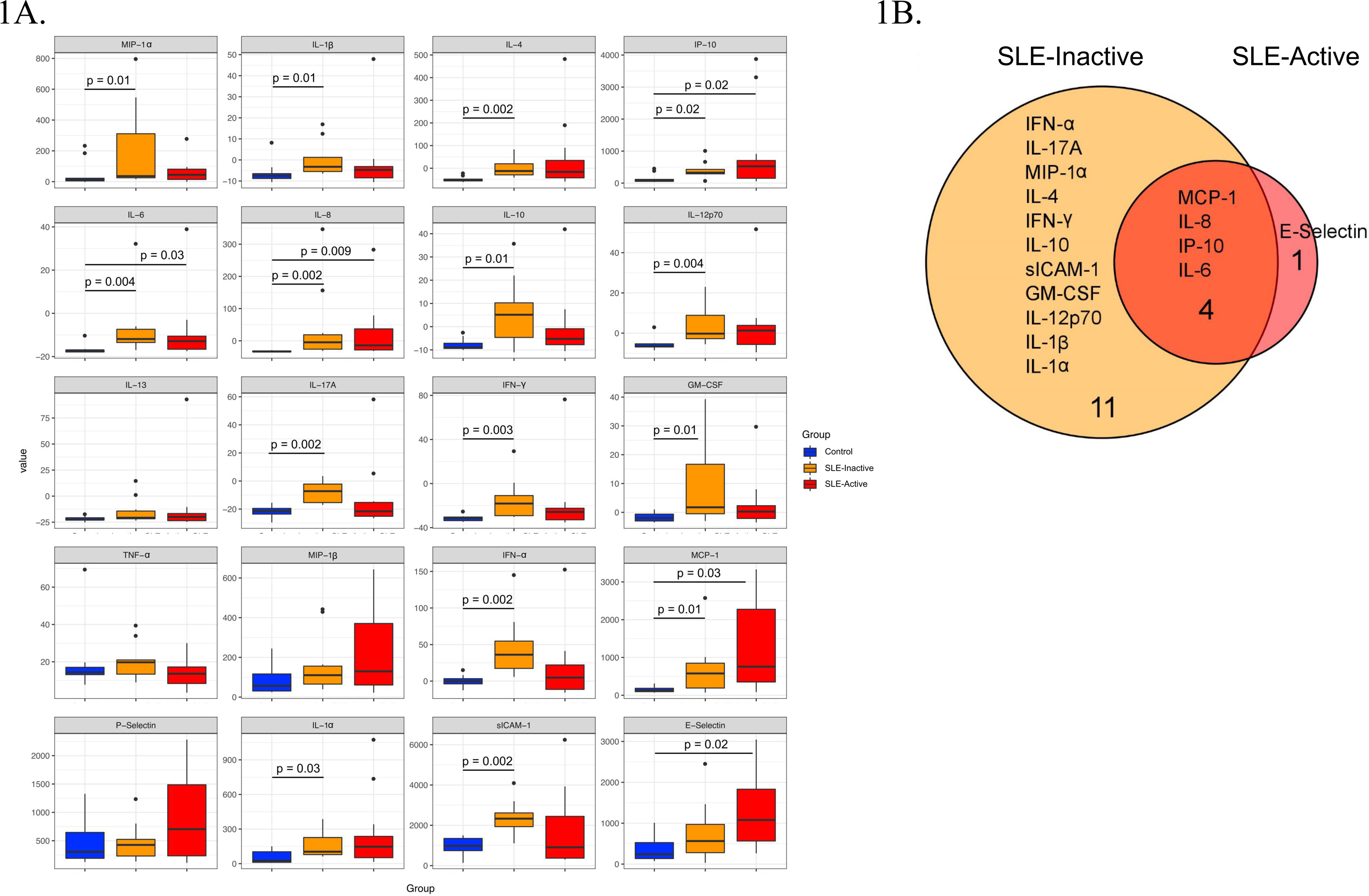
Human inflammation cytokines among control and SLE groups. (A) Variation in host cytokine response across control subjects and subjects with SLE-active and SLE-inactive conditions. The y-axis specifies the cytokine levels measured by a multiplex immuno-bead assay. Horizontal bar signifies the mean value for each group. Significance was evaluated by non-parametric Wilcoxon rank-sum test with Benjamini & Hochberg adjusted p-values. (B) A Venn diagram illustrating overlapping/non-overlapping cytokines in SLE-inactive and SLE-active conditions when compared to their respective controls. Venn diagrams are produced using VennDIS software (n = 91; p-values for two group comparisons are provided in Table 2).

**Table 2.** Beta-diversity pairwise comparisons between sample groups.

| Table 2. Beta-diversity pairwise comparisons between sample groups |  |  |  |  |  |  |
| --- | --- | --- | --- | --- | --- | --- |
| Factors | Groups | Non-periodontitis |  |  | Periodontitis |  |
|  |  | Healthy | SLE Active | SLE Inactive | Healthy | SLE Active |
| Non-periodontitis | SLE Active | 0.253 | - | - | - | - |
|  | SLE Inactive | 0.596 | 0.293 | - | - | - |
| Periodontitis | Healthy | 0.911 | 0.045* | 0.606 | - | - |
|  | SLE Active | 0.38 | 0.701 | 0.559 | 0.253 | - |
|  | SLE Inactive | 0.253 | 0.329 | 0.617 | 0.045* | 0.559 |

### 3.3 Subgingival Microbial Compositions in SLE

To visualize variations in the composition of subgingival biofilm-associated bacterial species across samples within SLE with or without periodontitis from controls (n = 31), SLE-inactive (n = 28), and SLE-active (n = 31) groups; we performed a collection of samples by probing the gingival site with PerioPaper strips, following elution. After hybridization with specific periodontal pathogen probes, DNA-checkerboard revealed pathogen signatures and abundances. A principal coordinate analysis (PCoA) on the Jaccard distances was generated taking into consideration of periodontal status stratification (Figure 2A). These visual patterns were confirmed by beta-diversity analysis using pairwise permutation Multivariate analysis of variance (MANOVA), revealing significant differences in bacterial species in the SLE-active group without periodontitis and SLE-inactive group with periodontitis compared to control group with periodontitis (p = 0.045; Table 2). Lastly, no differences were found between SLE-inactive groups and healthy controls (Supplementary Table 2). Taken together, we found significant variations in the microbiome composition in the SLE when compared to controls, with increased differences when SLE groups were stratified into SLE-A and SLE-I.

**Figure 2.**
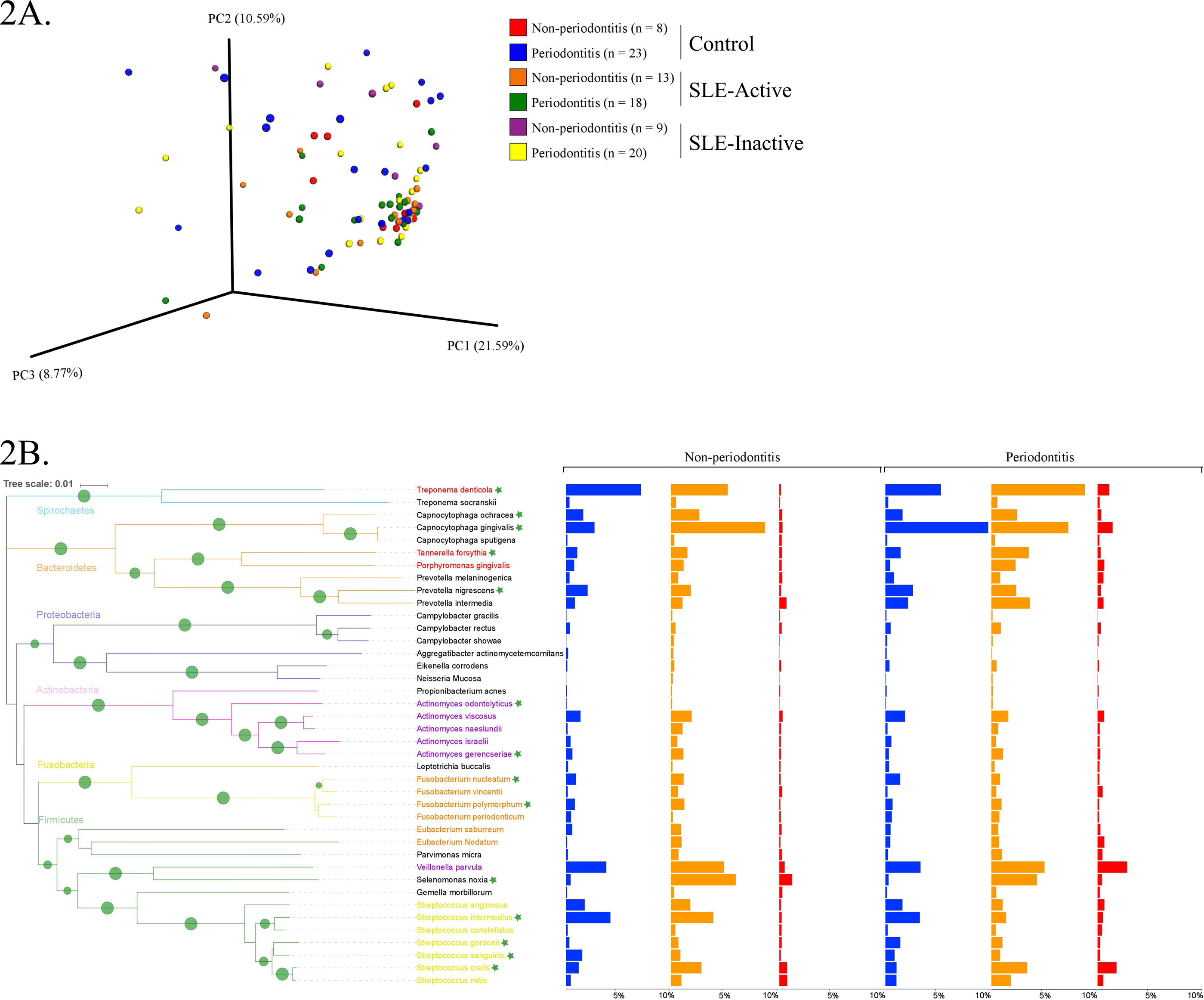
Bacterial composition in control and SLE individuals. (A) Principal coordinates analysis (PCoA) of the bacterial community between control and SLE individuals. The principal coordinates were calculated using Jaccard distance metric based on a binary matrix representing the presence/absence of the microbes across control and SLE subjects. Distance among samples in control versus inactive and active SLE conditions are visualized through EMPeror tool. The significance of separation between groups was tested by applying PERMANOVA test on the principal coordinates. (B) Prevalence of disease associated with bacterial species in non-periodontitis and periodontitis individuals. Bar graphs from left to right indicate differences in relative abundance of microbial species in control (in blue), SLE-inactive (in orange) and SLE-active individuals (in red). Branches in the tree are colored according to the phylum and periodontal pathogens are colored according to the red, purple, orange and yellow bacterial complex designations. Nodes with bootstraps higher than 80% are displayed with the tree. Indicator species with significant differences in abundances between periodontitis and non-periodontitis subjects are indicated with asterisks.

### 3.4 Microbial Signatures Associated with SLE and Periodontal Phenotypes

We next sought to understand the underlying microbial associations among SLE subjects and their clinical phenotypes. To investigate the subgingival plaque bacterial composition, we thus applied Wilcoxon rank-sum test on two datasets related to SLE phenotypes. The first dataset was based on relative abundance and the second dataset was based on logistic regression model binary abundances (presence/absence) to evaluate prevalence. Based on the relative abundance profiles, and the hierarchical clustering using Spearman’s rank correlation coefficients, we identified two distinct groups of bacterial species in SLE individuals with and without periodontitis when compared to their respective controls (Figure 2B, Figure 3A). Intriguingly, these comprised of species representatives of pathogenic groups previously classified in clusters including the red complex ^30^ (*Treponema denticola* and *Tannerella forsythia*), purple complex (*Actinomyces odontolyticus, A. gerencseriae*), orange complex (*Fusobacterium nucleatum*, *F. polymorphum*) and yellow complex (*Streptococcus intermedius*, *S. gordonii*, *S. sanguinis*, *S. oralis*). Additionally, *Capnocytophaga ochraceae*, *C. gingivalis*, *Prevotella nigrescens* and *Selenomonas noxia* showed significant differences in their relative abundances among SLE and control subjects.

**Figure 3.**
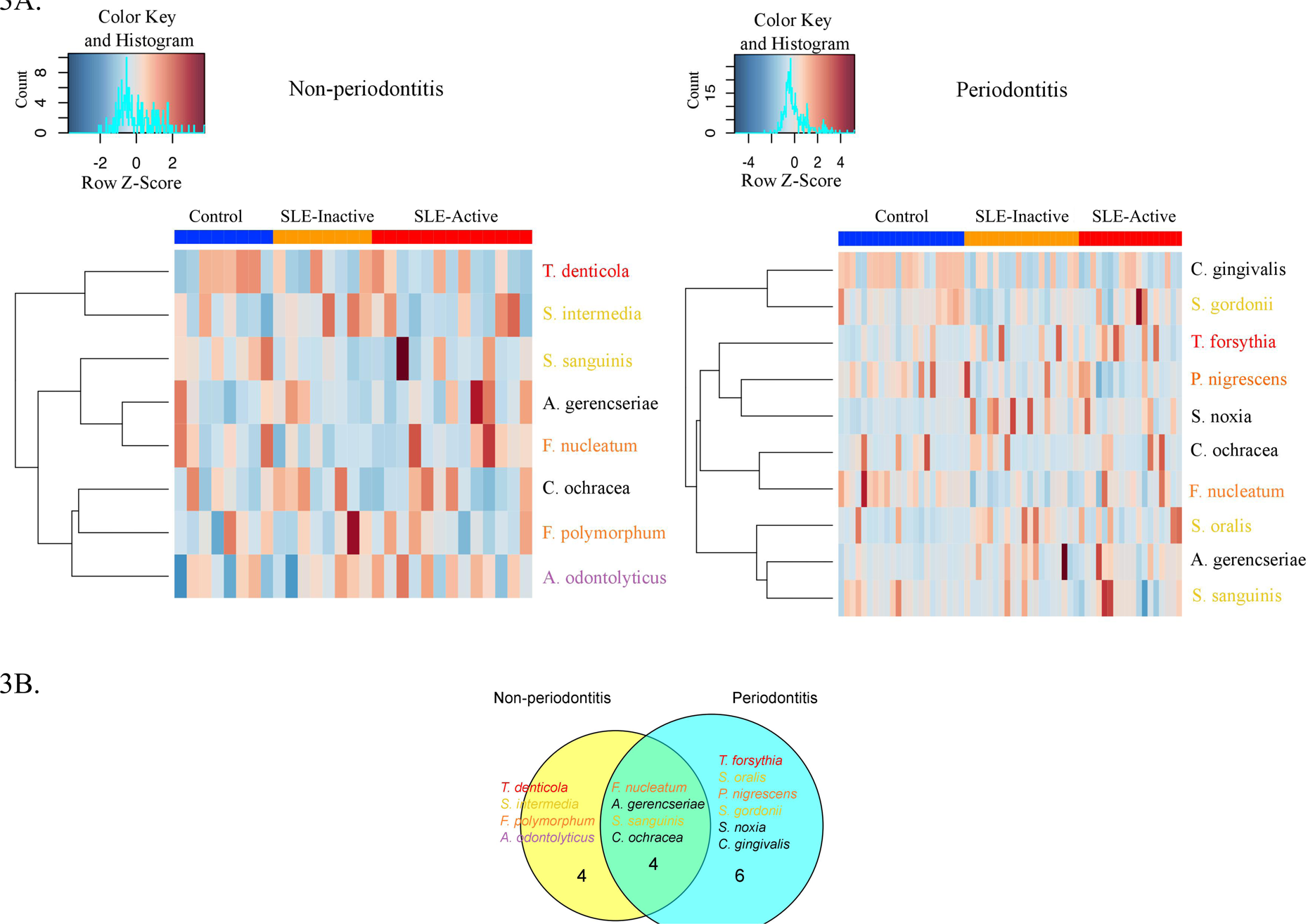
Non-periodontitis and periodontitis associated microbial species in SLE. (A) Heatmap illustrating the clustering of fourteen bacterial species with significant changes in abundances across control and SLE subjects. Subjects are shown in columns, while bacterial species shown in rows are colored by bacterial complex designations. A red and blue color affected by the row z-score signifies a higher and lower abundance of bacterial species, respectively. Significance was evaluated by Wilcoxon rank-sum test using Benjamini & Hochberg adjusted p-value < 0.05. (B) A Venn diagram illustrating shared and uniquely perturbed bacterial species across non-periodontitis and periodontitis individuals with SLE conditions. In each group, subjects were compared to their respective controls. The number of species differentially abundant in the comparison group. Venn diagrams are produced using VennDIS software.

When further stratifying the SLE comparisons according to periodontal status, the results showed that systemic comparisons were significantly different according to periodontal status (Supplementary Table 2). A total of twenty-four bacterial species had significant differential abundances among SLE and control subjects, with nine in the non-periodontitis and fifteen in the periodontitis groups (Figure 3A, and p-values listed in Supplementary Table 2). In non-periodontitis subjects, SLE-I versus healthy controls showed no significant microbial changes, demonstrating similar microbial abundances among these groups. In contrast, SLE-A versus healthy controls showed five significant microbial differences including *C. ochracea, F. nucleatum, S. sanguinis, A. gerencseriae, T. denticola*. When comparing SLE-A versus SLE-I subjects, four signatures were significant including, *C. ochracea, S. intermedia*, *A. odontolyticus, F. polymorphum.* Whereas in the periodontitis subjects, SLE-I versus healthy controls showed three significant microbial changes *A. gerencseriae, S. oralis, S. noxia.* An increase to seven significant microbial abundance changes were observed on SLE-A versus healthy controls comparison, including, *C. ochracea, F. nucleatum, P. nigrescens, S. sanguinis, C. gingivalis, T. forsythia, S. gordonii.* Finally, when comparing SLE-A versus SLE-I subjects, five signatures were significantly different including, *C. ochracea, P. nigrescens, A. gerencseriae, S. oralis, and T. forsythia.* While the long-term chronically inflamed SLE-I subjects showed higher abundances of periodontal pathogens when compared to SLE-A, SLE-I only showed significance difference to healthy controls in periodontitis groups. Overall the results suggest that the presence of periodontitis drove most of significant microbial differences among the SLE systemic status comparisons (SLE-I or SLE-A versus healthy controls, and SLE-A versus SLE-I).

We further characterized the periodontal pathogens in sites from SLE-I versus SLE-A individuals. Proportions of *C. gingivalis*, *S. gordonii*, *P. nigrescens*, *C. ochracea*, *F. nucleatum* and *S. sanguinis* were significantly reduced on SLE subjects (Figure 2B, Figure 3A, Supplementary Table 2). Based on our analysis the red complex pathogen, *T. forsythia*, was enriched in periodontitis subjects from SLE-active, but not in their healthy counterpart. In addition to these changes, we found *S. noxia*, *S. oralis* and *A. gerencseriae* at higher abundance in the SLE-I group compared with control individuals with periodontitis (Figure 2B, Figure 3A). *A. gerencseriae, S. oralis, C. ochracea, P. nigrescens* and *T. forsythia* were also found to be significantly different between SLE-I and SLE-A groups. These results suggest unique pathogen signatures associated with SLE-A phenotype and that commensals are associated with better systemic and periodontal health outcomes. A more stringent evaluation of bacterial abundance profiles via binary abundances revealed significant differences in the presence/absence of *C. ochracea, C. gingivalis, P. nigrescens, T. forsythia, F. nucleatum, S. gordonii* and *S. sanguinis* in periodontal sites of the SLE-active group compared with the control group (Supplementary Table 3).

Unique and overlapping microbial species were identified among periodontitis versus non-periodontitis groups (Figure 3B). Among fourteen unique microbial species, four bacterial species overlapped among non-periodontitis and periodontitis individuals (*F. nucleatum*, *A. gerencseriae*, *S. sanguinis* and *C. ochracea).* While four unique bacterial species (*T. denticola*, *S. intermedia*, *F. polymorphum* and *A. odontolyticus*) were significantly abundant only in non-periodontitis subjects, six species were unique in periodontitis subjects (*T. forsythia*, *S. oralis*, *P. nigrescence*, *S. gordonii*, *S. noxia* and *C. gingivalis*; Figure 3B). Together these results suggest periodontal bacterial diversity were lower in SLE patients with enrichment of specific periodontal pathogens dominating the microbial environment.

### 3.5 Bacterial Species and Cytokine Co-Occurrences

Periodontitis is caused by the complex interplay between subgingival microbiota composition and the host immune response. To unreveal functional insights, we investigated microbe-inflammatory cytokines correlations in periodontal and SLE phenotypes. The analysis showed associations between bacterial abundance and host cytokine levels in control, SLE-inactive and SLE-active individuals by Spearman’s rank-based correlations (Figure 4 A-C, Supplementary Figure 1). Among the relationships found, a total number of seventy-seven connecting edges of cytokine-cytokine, microbe-microbe, and microbe-cytokine was evident on SLE-A when compared to controls and SLE-I (Figure 4B and 4C). The controls showed negative values for most cytokines thus excluded from correlation plots. The majority of differentially abundant bacterial species perturbed cytokine levels, except *S. mitis*, which had more connectivity with other bacterial species in SLE individuals versus healthy controls.

**Figure 4.**
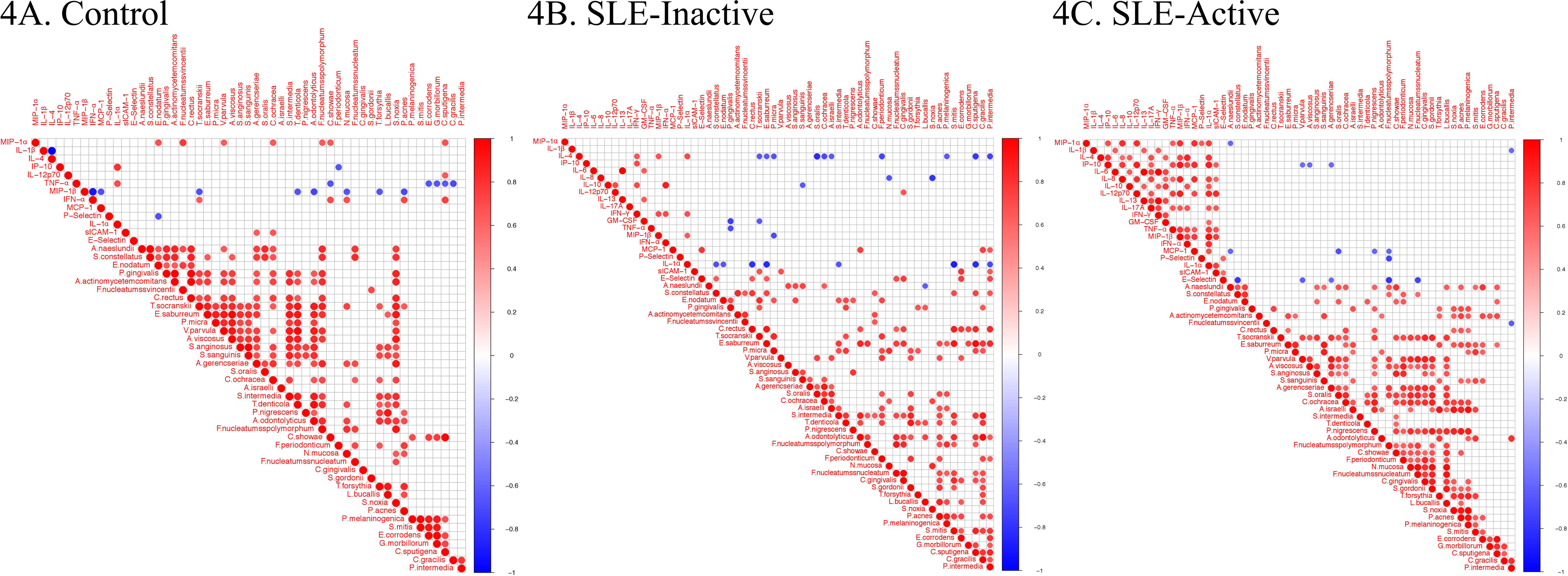
Bacterial species and cytokine co-occurrences. Spearman rank based pairwise correlation analysis between bacterial species and inflammatory cytokines (A) in control, (B) SLE-inactive and (C) SLE-active subjects. Only significant correlations (*p<0.05*) were plotted. The size of the spheres represents p-value. While strong correlations are shown by large circles, weak correlations are shown by small circles. The color of the circle denotes the strength of the correlation. Perfect positive correlation (with correlation coefficient 1) are indicated in dark red, whereas perfect inverse correlation (with correlation coefficient −1) are colored in dark blue.

In control individuals, the presence of pathogenic species, such as *T. denticola*, *A. odontolyticus* and *T. forsythia*, was found to be negatively correlated with MIP-1β levels (Figure 4A, Supplementary Figure 1A); *C. ochracea* abundance positively correlated with MIP-1-ɑ, sICAM-1 while *F. polymorphum* positively correlated with MIP-1-ɑ levels. *A. gerencseriae* is positively correlated with E-selectin levels. In SLE-inactive individuals, we found co-occurrence of *F. nucleatum* with E-selectin levels (Figure 4B, Supplementary Figure 1B). In contrast, the presence of health-related *S. sanguinis* inversely correlated with IP-10, while *A. odontolyticus* and *C. gingivalis* were positively associated with MCP-1 and IL12p70 respectively. *C. ochracea* inversely correlated with IL-4. *S. intermedia* inversely correlated with IL-1ɑ. *S. oralis* negatively correlated with IL-4. *S. noxia* negatively correlated with IL-8. In SLE-active individuals, *A. gerencseriae* is found to vary inversely with IP-10. *F. polymorphum* negatively correlated with MCP-1, E-selectin, and P-selectin. *S. oralis* and *P. nigrescens* showed negative associations with MCP-1 (Figure 4C, Supplementary Figure 1C).

The degree of connectivity (*i.e.* number of connecting edges to neighboring nodes) of *F. nucleatum* with other bacterial species and cytokines was higher in individuals with SLE when compared to control individuals. *A. gerencseriae, C. ochracea*, and *T. forsythia* showed higher connectivity in control and in SLE-A individuals while connectivity of *S. oralis* and *P. nigrescens* were elevated in SLE-A subjects. *S. noxia* was higher in the control group while *C. gingivalis* had no connectivity in control individuals (Figure 5). These findings suggest that subgingival bacterial species associated with SLE govern systemic host cytokine patterns impacting overall health.

**Figure 5.**
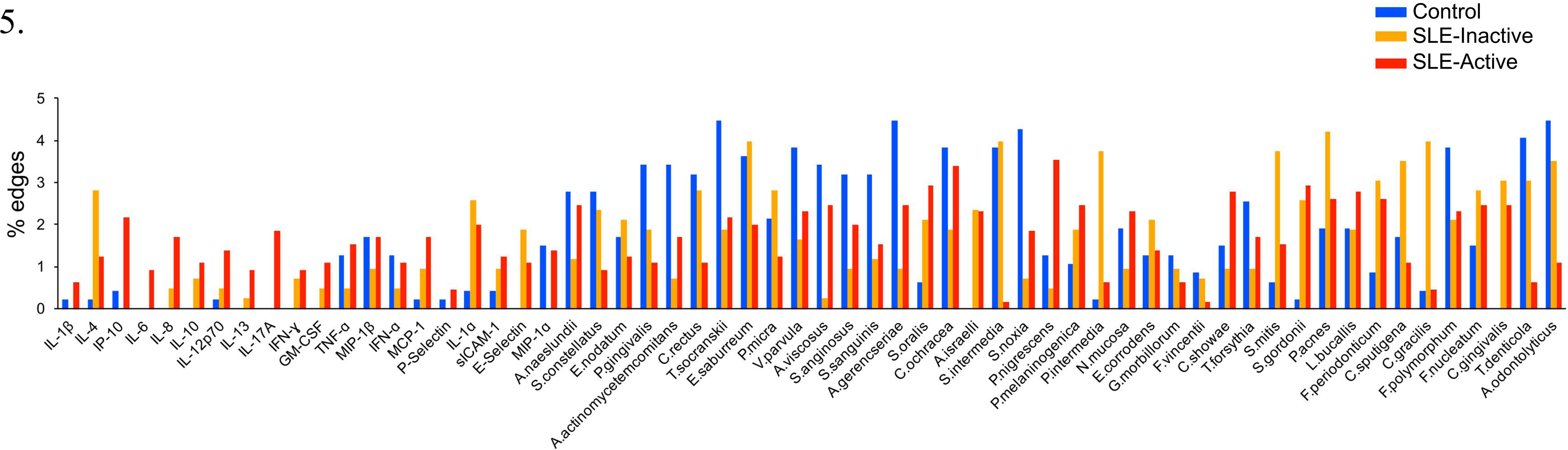
Topological analysis of the co-occurrence networks among cytokines and bacterial species. The selected topological parameter illustrates the percentage connectivity of each bacterial species and cytokines within control, SLE-I, and SLE-A networks.

## Discussion

At the center of autoimmune pathogenesis, this study provides evidence that inflammatory response to microbiome controls the severity, and magnitude of the SLE disease ^40^. The data also indicates that periodontal levels are also associated with low-grade systemic inflammation. Oral microbes, including subgingival bacteria, are involved in homeostasis and maintenance of health, and also in the initiation and progression of chronic periodontitis (CP) which leads to systemic inflammation such as SLE ^41–47^. Dual relationships among systemic and oral diseases play a role in the pathogenesis, and low-grade inflammation modulating the biological compartments ^19^, and here we assayed how the microbial species influence specific systemic cytokines. Evidence of the exact repertoire of subgingival pathogens influencing specific lupus phenotypes has been scarcer in the context of oral and systemic disease. We explored two distinct populations, healthy and SLE positive patients to determine these correlations. Our clinical cross-sectional study is unique because we have stratified chronic SLE patients, which are usually under treatment, (SLE-inactive) and a more acute group, and recently diagnosed SLE subjects (SLE-active). We have also carefully stratified the subjects in periodontally compromised versus healthy controls. Our results suggest that chronic and SLE-inactive subjects are positively associated with severe periodontitis states. Among healthy and SLE groups, age was not significantly different, but when stratifying SLE active and inactive groups, younger subjects were found significantly present on SLE-A when compared to SLE-I. This was expected due to time of disease since diagnosis ^48^. Patients with SLE in general showed clear molecular patterns related to chronic inflammation and immune response to microbiome when compared to healthy controls.

A more distinct host cytokine dysregulation was associated with SLE subjects when compared to healthy controls. The data presented herein show no evidence that SLE-active subjects had higher serum concentrations of pro- and anti-inflammatory cytokine. But, both SLE groups showed higher expressions than healthy control. Interestingly IL-10 which is a anti-inflammatory cytokine response was found increased in SLE-I inactive when compared to active and controls (p=0.01, Figure 1A). Our results showed significantly increased levels of IFN-ɑ, IL-17A, MIP-1-ɑ, IL-4, IFN-ɣ, and IL-10 in SLE-inactive individuals. sICAM-1, GM-CSF, IL-12p70, IL-1β, IL-1ɑ in serum were unique of SLE-inactive subjects, while four cytokines were significantly increased, with one unique profile including E-selectin of SLE-A subjects (Figure 1B). A high number of cytokine perturbations in SLE-I individuals can be associated with diverse clinical manifestations, activity and the severity of the disease. As mediators of inflammation, cytokine production feeds forward cell-cell and cell-tissue communications guiding health and disease phenotypes and organ disruptions. Here, we show that proinflammatory cytokines MCP-1, IL-8, IP-10, and IL-6 were significantly elevated in SLE-I and SLE-A patients in comparison to non-SLE subjects (Figure 1A). Of this proinflammatory cytokines, IL-6, IL-17, TNF-α, and IFN-α suppression have been reported as therapeutic targets for clinical management of lupus and other chronic inflammatory diseases^4,49–51.^

Systemic and local periodontal inflammation can impact subgingival bacterial compositions, which can in turn further enhance systemic inflammation, leading to tissue loss, mainly in subjects with SLE condition ^49^. The bacterial abundances of fourty-selected subgingival species were evaluated through checkerboard and we found that two distinct groups of subgingival bacteria were present in SLE individuals, especially when stratifying for the presence of absence of periodontitis (Figure 3A). *S. sanguinis* and *C. ochracea* are known to be prevalent in healthy subgingival sites ^52–54^. In line with previous studies, proportions of *T. denticola*, *S. sanguinis* and *C. ochracea* were significantly higher in periodontally healthy individuals than in individuals with SLE. *A. gerencseriae*, and *F. nucleatum* were more evidently abundant in SLE individuals without periodontitis. We found a higher abundance of *S. noxia*, *S. oralis* and *A. gerencseriae* in SLE subjects with periodontitis. On the other hand, we found significant depletion of *C. gingivalis*, *S. gordonii*, *P. nigrescens*, *C. ochracea*, *F. nucleatum* and *S. sanguinis*, which were previously found to be associated with SLE pathogenesis (Figure 3A, Supplementary Table 2, Supplementary Table 3). It is noteworthy that SLE-I microbial compositions was not different from healthy controls in the non-periodontitis group, indicating that they are similar in periodontally healthy subjects (Supplementary Table 2). On the contrary, we show that SLE-A microbial positions maintains differences in both periodontitis and non-periodontitis groups, and that one form of inflammation (local or systemic) control these differences associated with SLE systemic inflammatory profiles.

Co-occurrence of subgingival bacteria and cytokines also differed among controls, SLE-I and SLE-A subjects. Co-occurrence among cytokines was more evident (77 positive connecting edges) in the SLE states, suggesting positive association among cytokines are an important driver of SLE pathogenesis (Figure 4A-C). We did not find any correlations between bacteria and increased levels of IL-6 and IL-17. Despite IFN-α and TNF-α co-occurred with certain subgingival bacteria, these bacteria were not differentially abundant. Beyond cytokine to microbe interactions, SLE-A showed higher cytokine-cytokine co-occurrences. IL-1β, IL-8, and IL-12 are associated with increased innate and adaptive immune response including T-cell. IL-1β with high inflammatory properties has been shown to increase in SLE condition with systemic manifestations, especially in the inflammasome activation associated with inflammatory responses to extracellular pathogens. IL-8 has been shown to induce neutrophil recruitment, especially during early stages of SLE. Qiu *et al*. reported high levels of IL-12 family-related cytokines in triggering the production of anti-double-stranded DNA antibodies ^55^.

Functional microbiome derived inflammation data at the host and microbial levels are necessary to make assumptions on ecological systems from the oral cavity and its influences on systemic disease phenotypes. The data presented here generate additional questions about the extent to which detrimental and protective microbiome features are innately acquired and significantly modulated by the host systemic conditions. Overall the results offer a new perspective on the host-microbial interactions in lupus disease and periodontal pathogenic environments. Consequently, these results favor the hypothesis of oral-systemic inflammation linking both conditions. Moreover, the realization that there remain high anti-inflammatory profiles on subjects with inactive disease when compared to active, offer a mechanistic hypothesis to the lack of disease comorbidities of this suppressed condition, but the host is still able to produce low-grade inflammation systemically. In future studies using next generation sequencing strategies of highly diverse SLE subjects, it will be possible to map the microbial ecology of multiple sites of the human body, including oral sites. As the science of host-microbes is advancing, highly individualized, rather than simple binary distinctions of healthy and disease or single-microbe etiology across populations, might be found ^56^.

The results presented here show that oral-systemic pathogenic burden is evident in lupus subjects. These data are also consistent with complex ecological interactions among multiple human chronic diseases ^57^. Among the relationships found, co-occurrence within the cytokines was evident on SLE-A group (a total number of 77 connecting edges, Figure 4C) when compared to controls and SLE-inactive states (Figure 4A and Figure 4B), suggesting positive relationships of proinflammatory cytokines as important drivers of SLE pathogenesis. Currently, there is no cure for SLE manifestations, and administration of immunosuppressive drugs are aimed at suppressing immune signals chronically, and removal of the etiological factors nor disruption of the human microbiome. It is possible that by controlling oral microbiome-induced inflammation, the systemic low-grade inflammation burden will be reduced, preventing future emergence of comorbidities and disease clusters. Thus, periodontal and oral diseases are biological sources of complex ecosystem influencing systemic inflammation. These findings, along with future longitudinal studies, could provide improved resolution on how microbial and immune features interact to modify disease outcomes on selected individuals and populations. This strategy is key to trace how health and disease-associated oral biomarkers are spatially and temporally regulated in the context of chronic human diseases.

## Abbreviations

AAP: American Academy of Periodontology
ACR: American College of Rheumatology
ANA: Abnormal antinuclear antibodies
BOP: Bleeding on probing
CAL: Clinical attachment loss
CDC: Centers for Disease Control and Prevention
CP: Chronic periodontitis
E-selectin: Endothelial-selectin
GCF: Gingival crevicular fluid
GM-CSF: Granulocyte-macrophage colony-stimulating factor
HOMD: Human Oral Microbiome Database
IFN-ɑ: Interferon alpha
IFN-ɣ: Interferon gamma
IL-10: Interleukin-10
IL-12p70: Interleukin-12p70
IL-13: Interleukin-13
IL-17A: Interleukin-17A
IL-1ɑ: Interleukin-1 alpha
IL-1β: Interleukin-1 beta
IL-4: Interleukin-4
IL-6: Interleukin-6
IL-8: Interleukin-8
IP-10: Interferon gamma-induced protein 10
TNF-ɑ: Tumor necrosis factor alpha
iTOL: Interactive tree of life
MCP-1: Monocyte chemoattractant protein 1
P-Selectin: Platelet-selectin
sICAM-1: Soluble intercellular adhesion molecule-1
MEGA: Molecular Evolutionary Genetics Analysis software
MIP-1-ɑ: Macrophage inflammatory protein 1 alpha
MIP-1β: Monocyte/macrophage inflammatory protein-1 beta
PBS: Phosphate Buffer Saline
PCoA: Principal Coordinates Analysis
PD: Probing depth
SLE: Systemic lupus erythematosus
SLE-A: SLE-active
SLE-I: SLE-inactive
SLEDAI: Systemic lupus erythematosus disease activity index
SLICC/ACR-DI: Systemic Lupus International Collaborating Clinics of American College of Rheumatology Damage Index
SLICC: Systemic lupus international collaborating clinics
FDR: False discovery rate

## Supplementary Information

**Supplementary Table 1.** Comparison of cytokine levels between SLE and control groups. Significance was evaluated by the non-parametric Wilcoxon rank-sum test. Significant p-values (Benjamini & Hochberg adjusted) are highlighted in bold.

**Supplementary Table 2.** Comparison of relative abundances of bacterial species between SLE and control groups with or without periodontitis. Significance was evaluated by the non-parametric Wilcoxon rank-sum test. Significant p-values (Benjamini & Hochberg adjusted) are highlighted in bold.

**Supplementary Table 3.** Comparison of presence/absence of bacterial species between SLE and control groups with or without periodontitis. Significance was evaluated by the logistic regression model. Significant p-values (Benjamini & Hochberg adjusted) are highlighted in bold.

**Supplementary Table 4.** Comprehensive demographics table and clinical comparisons of SLE and control subjects with chronic or non-chronic periodontitis..

**Supplementary Figure 1.**
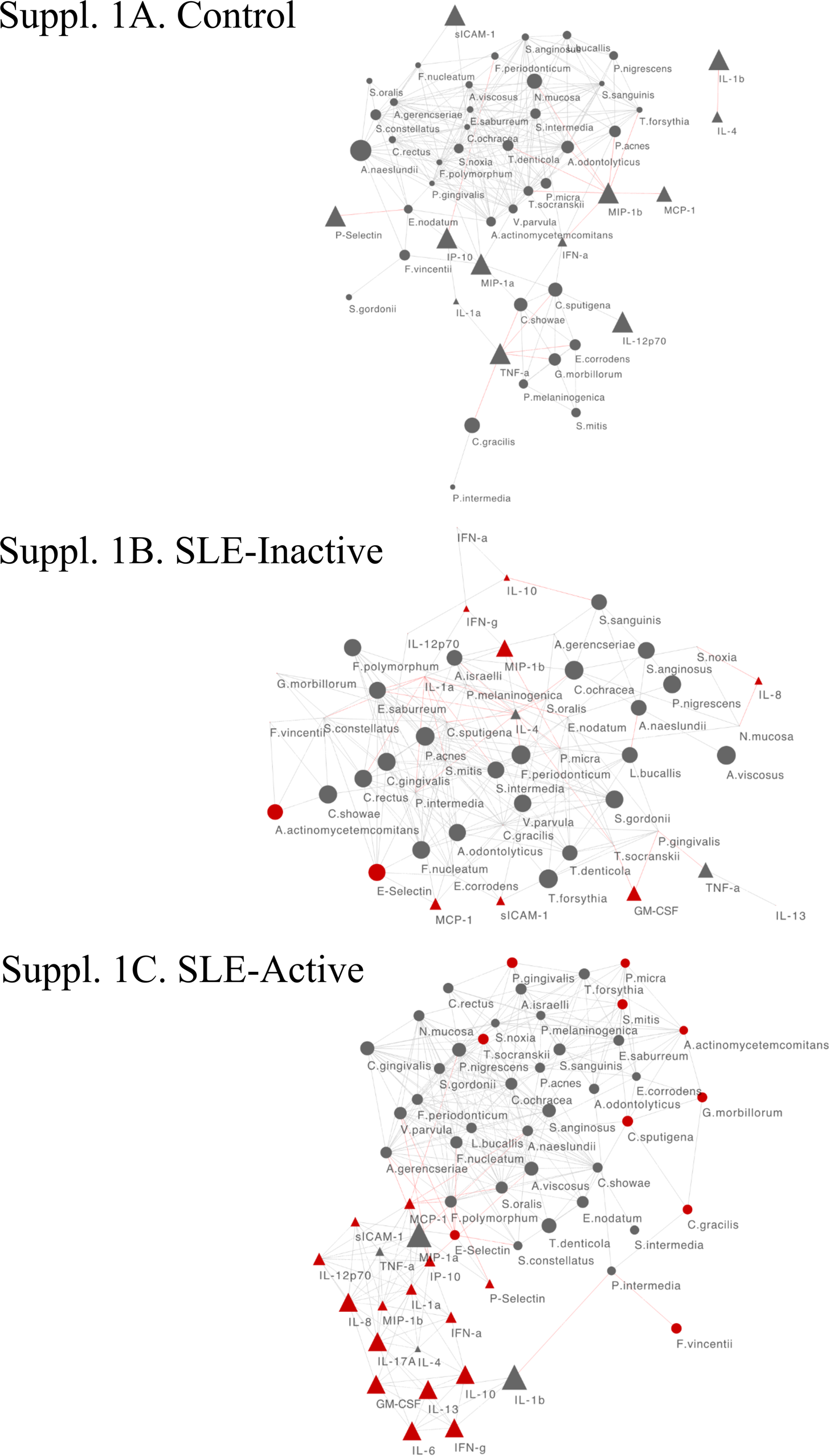
Significant Spearman correlations among bacteria and host cytokines in (A) control, (B) SLE-inactive, and (C) SLE-active states. The size of the node (circles) in SLE-inactive and the SLE-active group represents a log2 fold change of abundances in comparison to control subjects. Increase in the size of the node signifies lower expression in control when compared to SLE subjects. Edges in red represent negative correlations, gray represent positive correlations.

## Acknowledgements

This work was performed at the Forsyth Institute, J. Craig Venter Institute and at the University of Brasilia. This work was supported in parts by the National Institutes of Health [DE023584. 2018]; and the J. Craig Venter Innovation Fund given to MF. There are no conflicts of interest with any of the authors.

## References

1. Scher JU, Bretz WA, Abramson SB. Periodontal disease and subgingival microbiota as contributors for rheumatoid arthritis pathogenesis: modifiable risk factors? Curr Opin Rheumatol. 2014 Jul;26(4):424–429. PMCID: PMC4128331

2. Kobayashi T, Ito S, Yasuda K, Kuroda T, Yamamoto K, Sugita N, Tai H, Narita I, Gejyo F, Yoshie H. The Combined Genotypes of Stimulatory and Inhibitory Fcγ Receptors Associated With Systemic Lupus Erythematosus and Periodontitis in Japanese Adults. J Periodontol. 2007 Mar;78(3):467–474.

3. Corrêa JD, Saraiva AM, Queiroz-Junior CM, Madeira MFM, Duarte PM, Teixeira MM, Souza DG, da Silva TA. Arthritis-induced alveolar bone loss is associated with changes in the composition of oral microbiota. Anaerobe. 2016 Jun;39:91–96. PMID: 26996070

4. Corrêa JD, Calderaro DC, Ferreira GA, Mendonça SMS, Fernandes GR, Xiao E, Teixeira AL, Leys EJ, Graves DT, Silva TA. Subgingival microbiota dysbiosis in systemic lupus erythematosus: association with periodontal status. Microbiome. 2017 Mar 20;5(1):34. PMCID: PMC5359961

5. Socransky SS, Haffajee AD, Cugini MA, Smith C, Kent RL. Microbial complexes in subgingival plaque [Internet]. Journal of Clinical Periodontology. 1998. p. 134–144. Available from: http://dx.doi.org/10.1111/j.1600-051x.1998.tb02419.x

6. Kumar PS, Leys EJ, Bryk JM, Martinez FJ, Moeschberger ML, Griffen AL. Changes in periodontal health status are associated with bacterial community shifts as assessed by quantitative 16S cloning and sequencing. J Clin Microbiol. 2006 Oct;44(10):3665–3673. PMCID: PMC1594761

7. Armitage GC. Development of a Classification System for Periodontal Diseases and Conditions [Internet]. Annals of Periodontology. 1999. p. 1–6. Available from: http://dx.doi.org/10.1902/annals.1999.4.1.1

8. Rees F, Doherty M, Grainge MJ, Lanyon P, Zhang W. The worldwide incidence and prevalence of systemic lupus erythematosus: a systematic review of epidemiological studies. Rheumatology. 2017 Nov 1;56(11):1945–1961. PMID: 28968809

9. Roper G. Lupus Awareness Survey for the Lupus Foundation of America [Executive Summary Report]. Washington, DC GfK Roper Public Affairs & Corporate Communications. 2012;

10. Pons-Estel GJ, Alarcón GS, Scofield L, Reinlib L, Cooper GS. Understanding the epidemiology and progression of systemic lupus erythematosus. Semin Arthritis Rheum. 2010 Feb;39(4):257–268. PMCID: PMC2813992

11. Wallace D, Hahn BH. Dubois’ Lupus Erythematosus and Related Syndromes E-Book: Expert Consult – Online and Print. Elsevier Health Sciences; 2012.

12. Hochberg MC. Updating the American College of Rheumatology revised criteria for the classification of systemic lupus erythematosus. Arthritis Rheum. 1997 Sep;40(9):1725. PMID: 9324032

13. Agmon-Levin N, Mosca M, Petri M, Shoenfeld Y. Systemic lupus erythematosus one disease or many? Autoimmun Rev. 2012 Jun;11(8):593–595. PMID: 22041578

14. Proal AD, Albert PJ, Marshall TG. The human microbiome and autoimmunity. Curr Opin Rheumatol. 2013 Mar;25(2):234–240. PMID: 23370376

15. Azzouz D, Omarbekova A, Heguy A, Schwudke D, Gisch N, Rovin BH, Caricchio R, Buyon JP, Alekseyenko AV, Silverman GJ. Lupus nephritis is linked to disease-activity associated expansions and immunity to a gut commensal. Ann Rheum Dis [Internet]. 2019 Feb 19; Available from: http://dx.doi.org/10.1136/annrheumdis-2018-214856 PMID: 30782585

16. Mutlu S, Richards A, Maddison P, Scully C. Gingival and periodontal health in systemic lupus erythematosus. Community Dent Oral Epidemiol. 1993 Jun;21(3):158–161. PMID: 8348790

17. de Araújo Navas EAF, Sato EI, Pereira DFA, Back-Brito GN, Ishikawa JA, Jorge AOC, Brighenti FL, Koga-Ito CY. Oral microbial colonization in patients with systemic lupus erythematous: correlation with treatment and disease activity. Lupus. 2012 Aug;21(9):969–977. PMID: 22453994

18. Novo E, Garcia-MacGregor E, Viera N, Chaparro N, Crozzoli Y. Periodontitis and Anti-Neutrophil Cytoplasmic Antibodies in Systemic Lupus Erythematosus and Rheumatoid Arthritis: A Comparative Study [Internet]. Journal of Periodontology. 1999. p. 185–188. Available from: http://dx.doi.org/10.1902/jop.1999.70.2.185

19. van der Meulen TA, Harmsen HJM, Vila AV, Kurilshikov A, Liefers SC, Zhernakova A, Fu J, Wijmenga C, Weersma RK, de Leeuw K, Bootsma H, Spijkervet FKL, Vissink A, Kroese FGM. Shared gut, but distinct oral microbiota composition in primary Sjögren’s syndrome and systemic lupus erythematosus. J Autoimmun. 2019 Feb;97:77–87. PMID: 30416033

20. Petri M, Orbai A-M, Alarcón GS, Gordon C, Merrill JT, Fortin PR, Bruce IN, Isenberg D, Wallace DJ, Nived O, Sturfelt G, Ramsey-Goldman R, Bae S-C, Hanly JG, Sánchez-Guerrero J, Clarke A, Aranow C, Manzi S, Urowitz M, Gladman D, Kalunian K, Costner M, Werth VP, Zoma A, Bernatsky S, Ruiz-Irastorza G, Khamashta MA, Jacobsen S, Buyon JP, Maddison P, Dooley MA, van Vollenhoven RF, Ginzler E, Stoll T, Peschken C, Jorizzo JL, Callen JP, Lim SS, Fessler BJ, Inanc M, Kamen DL, Rahman A, Steinsson K, Franks AG Jr, Sigler L, Hameed S, Fang H, Pham N, Brey R, Weisman MH, McGwin G Jr, Magder LS. Derivation and validation of the Systemic Lupus International Collaborating Clinics classification criteria for systemic lupus erythematosus. Arthritis Rheum. 2012 Aug;64(8):2677–2686. PMCID: PMC3409311

21. Tan EM, Cohen AS, Fries JF, Masi AT, McShane DJ, Rothfield NF, Schaller JG, Talal N, Winchester RJ. The 1982 revised criteria for the classification of systemic lupus erythematosus. Arthritis Rheum. 1982 Nov;25(11):1271–1277. PMID: 7138600

22. Bombardier C, Gladman DD, Urowitz MB, Caron D, Chang CH, Austin A, Bell A, Bloch DA, Corey PN, Decker JL, Esdaile J, Fries JF, Ginzler EM, Goldsmith CH, Hochberg MC, Jones JV, Riche NGHL, Liang MH, Lockshin MD, Muenz LR, Sackett DL, Schur PH. Derivation of the sledai. A disease activity index for lupus patients. Arthritis & Rheumatism. 1992 Jun;35(6):630–640.

23. Gladman D, Ginzler E, Goldsmith C, Fortin P, Liang M, Sanchez-Guerrero J, Urowitz M, Bacon P, Bombardieri S, Hanly J, Jones J, Hay E, Symmons D, Isenberg D, Kalunion K, Maddison P, Nived O, Sturfelt G, Petri M, Richter M, Snaith M, Zoma A. The development and initial validation of the systemic lupus international collaborating clinics/American college of rheumatology damage index for systemic lupus erythematosus [Internet]. Arthritis & Rheumatism. 1996. p. 363–369. Available from: http://dx.doi.org/10.1002/art.1780390303

24. Ainamo J, Bay I. Problems and proposals for recording gingivitis and plaque. Int Dent J. 1975 Dec;25(4):229–235. PMID: 1058834

25. O’Leary TJ, Drake RB, Naylor JE. The Plaque Control Record [Internet]. Journal of Periodontology. 1972. p. 38–38. Available from: http://dx.doi.org/10.1902/jop.1972.43.1.38

26. Enikov ET, Eke E. Teaching Classical Control System Course With Portable Student-Owned Mechatronic Kits [Internet]. Volume 5: Education and Globalization; General Topics. 2012. Available from: http://dx.doi.org/10.1115/imece2012-86700

27. Eke PI, Dye BA, Wei L, Slade GD, Thornton-Evans GO, Borgnakke WS, Taylor GW, Page RC, Beck JD, Genco RJ. Update on Prevalence of Periodontitis in Adults in the United States: NHANES 2009 to 2012. J Periodontol. 2015 May;86(5):611–622. PMCID: PMC4460825

28. Feinberg AP, Vogelstein B. A technique for radiolabeling DNA restriction endonuclease fragments to high specific activity [Internet]. Analytical Biochemistry. 1983. p. 6–13. Available from: http://dx.doi.org/10.1016/0003-2697(83)90418-9

29. Brito LCN, Sobrinho APR, Teles RP, Socransky SS, Haffajee AD, Vieira LQ, Teles FRF. “Microbiologic profile of endodontic infections from HIV− and HIV+ patients using MDA and Checkerboard.” Oral Dis. NIH Public Access; 2012 Sep;18(6):558. PMCID: PMC4148015

30. Socransky SS, Haffajee AD, Smith C, Martin L, Haffajee JA, Uzel NG, Goodson JM. Use of checkerboard DNA-DNA hybridization to study complex microbial ecosystems [Internet]. Oral Microbiology and Immunology. 2004. p. 352–362. Available from: http://dx.doi.org/10.1111/j.1399-302x.2004.00168.x

31. Brito LCN, Sobrinho APR, Teles RP, Socransky SS, Haffajee AD, Vieira LQ, Teles FRF. Microbiologic profile of endodontic infections from HIV- and HIV+ patients using Multiple-Displacement Amplification and Checkerboard DNA-DNA Hybridization. Oral Dis. Wiley Online Library; 2012;18(6):558–567.

32. Vázquez-Baeza Y, Pirrung M, Gonzalez A, Knight R. EMPeror: a tool for visualizing high-throughput microbial community data. Gigascience. 2013 Nov 26;2(1):16. PMCID: PMC4076506

33. Kumar S, Stecher G, Tamura K. MEGA7: Molecular Evolutionary Genetics Analysis Version 7.0 for Bigger Datasets. Mol Biol Evol. 2016 Jul;33(7):1870–1874. PMID: 27004904

34. Letunic I, Bork P. Interactive tree of life (iTOL) v3: an online tool for the display and annotation of phylogenetic and other trees. Nucleic Acids Res. 2016 Jul 8;44(W1):W242–5. PMCID: PMC4987883

35. Chalan P, Bijzet J, van den Berg A, Kluiver J, Kroesen B-J, Boots AMH, Brouwer E. Analysis of serum immune markers in seropositive and seronegative rheumatoid arthritis and in high-risk seropositive arthralgia patients. Sci Rep [Internet]. Nature Publishing Group; 2016 [cited 2019 Apr 16];6. Available from: https://www.ncbi.nlm.nih.gov/pmc/articles/PMC4870704/ PMCID: PMC4870704

36. Lee CY, Robinson DA, Johnson CA Jr, Zhang Y, Wong J, Joshi DJ, Wu T-T, Knight PA. A Randomized Controlled Trial of Liposomal Bupivacaine Parasternal Intercostal Block for Sternotomy. Ann Thorac Surg. 2019 Jan;107(1):128–134. PMID: 30170012

37. Mosca M, Merrill JT, Bombardieri S. Assessment of disease activity in systemic lupus erythematosus. Systemic lupus erythematosus Philadelphia (PA): Elsevier. 2007;19–23.

38. Brugos B, Vincze Z, Sipka S, Szegedi G, Zeher M. Serum and urinary cytokine levels of SLE patients. Pharmazie. 2012 May;67(5):411–413. PMID: 22764573

39. McCarthy EM, Smith S, Lee RZ, Cunnane G, Doran MF, Donnelly S, Howard D, O’Connell P, Kearns G, Ní Gabhann J, Jefferies CA. The association of cytokines with disease activity and damage scores in systemic lupus erythematosus patients. Rheumatology. 2014 Sep;53(9):1586–1594. PMID: 24706988

40. Fava A, Petri M. Systemic lupus erythematosus: Diagnosis and clinical management. J Autoimmun. 2019 Jan;96:1–13. PMCID: PMC6310637

41. Cunningham LL, Novak MJ, Madsen M, Abadi B, Ebersole JL. A bidirectional relationship of oral-systemic responses: observations of systemic host responses in patients after full-mouth extractions [Internet]. Oral Surgery, Oral Medicine, Oral Pathology and Oral Radiology. 2014. p. 435–444. Available from: http://dx.doi.org/10.1016/j.oooo.2013.11.502

42. Glick M. The Oral-systemic Health Connection: A Guide to Patient Care. Quintessence Publishing Company; 2014.

43. T.j.c, T. JC. Oral diagnosis of systemic disease [Internet]. Oral Surgery, Oral Medicine, Oral Pathology. 1960. p. 896. Available from: http://dx.doi.org/10.1016/0030-4220(60)90028-1

44. Valentich MA, Cafaro TA, Serra HM. Oral Cavity-Associated Immune System: What is New? [Internet]. Current Immunology Reviews. 2011. p. 253–263. Available from: http://dx.doi.org/10.2174/157339511796196610

45. Olsen I, Yamazaki K. Can oral bacteria affect the microbiome of the gut? J Oral Microbiol. 2019 Mar 18;11(1):1586422. PMCID: PMC6427756

46. Kato T, Yamazaki K, Nakajima M, Date Y, Kikuchi J, Hase K, Ohno H, Yamazaki K. Oral Administration of Porphyromonas gingivalis Alters the Gut Microbiome and Serum Metabolome. mSphere [Internet]. 2018 Oct 17;3(5). Available from: http://dx.doi.org/10.1128/mSphere.00460-18 PMCID: PMC6193602

47. Kumar PS. From focal sepsis to periodontal medicine: a century of exploring the role of the oral microbiome in systemic disease [Internet]. The Journal of Physiology. 2017. p. 465–476. Available from: http://dx.doi.org/10.1113/jp272427

48. Fabbri C, Fuller R, Bonfá E, Guedes LKN, D’Alleva PSR, Borba EF. Periodontitis treatment improves systemic lupus erythematosus response to immunosuppressive therapy. Clin Rheumatol. 2014 Apr;33(4):505–509. PMID: 24415114

49. Jacob N, Stohl W. Cytokine disturbances in systemic lupus erythematosus. Arthritis Res Ther. 2011 Jul 6;13(4):228. PMCID: PMC3239336

50. Tackey E, Lipsky PE, Illei GG. Rationale for interleukin-6 blockade in systemic lupus erythematosus. Lupus. 2004;13(5):339–343. PMCID: PMC2014821

51. Nalbandian A, Crispín JC, Tsokos GC. Interleukin-17 and systemic lupus erythematosus: current concepts [Internet]. Clinical & Experimental Immunology. 2009. p. 209–215. Available from: http://dx.doi.org/10.1111/j.1365-2249.2009.03944.x

52. Galimanas V, Hall M, Singh N, Lynch MD, Goldberg M, Tenenbaum H, Cvitkovitch D, Neufeld J, Senadheera D. Bacterial community composition of chronic periodontitis and novel oral sampling sites for detecting disease indicators [Internet]. Microbiome. 2014. p. 32. Available from: http://dx.doi.org/10.1186/2049-2618-2-32

53. Wang J, Qi J, Zhao H, He S, Zhang Y, Wei S, Zhao F. Metagenomic sequencing reveals microbiota and its functional potential associated with periodontal disease. Sci Rep. 2013;3:1843. PMCID: PMC3654486

54. Wade WG. Characterisation of the human oral microbiome [Internet]. Journal of Oral Biosciences. 2013. p. 143–148. Available from: http://dx.doi.org/10.1016/j.job.2013.06.001

55. Qiu F, Song L, Yang N, Li X. Glucocorticoid downregulates expression of IL-12 family cytokines in systemic lupus erythematosus patients. Lupus. 2013 Sep;22(10):1011–1016. PMID: 23884985

56. Gomez A, Espinoza JL, Harkins DM, Leong P, Saffery R, Bockmann M, Torralba M, Kuelbs C, Kodukula R, Inman J, Hughes T, Craig JM, Highlander SK, Jones MB, Dupont CL, Nelson KE. Host Genetic Control of the Oral Microbiome in Health and Disease. Cell Host Microbe. 2017 Sep 13;22(3):269–278.e3. PMCID: PMC5733791

57. Blaser MJ. Who are we? Indigenous microbes and the ecology of human diseases. EMBO Rep. 2006 Oct;7(10):956–960. PMCID: PMC1618379

