## Supplementary Table 1 for "Host-Microbial Interactions in Systemic Lupus Erythematosus and Periodontitis"

| Cytokine | SLE-I vs Control | SLE-A vs Control | SLE-A vs SLE-I |
|---|---|---|---|
| IL-17A | **0.002298116** | 0.79174097 | 0.2713651 |
| IFN-a | **0.002298116** | 0.65350477 | 0.2713651 |
| sICAM-1 | **0.002298116** | 0.64712721 | 0.5034782 |
| E-Selectin | 0.241736248 | **0.02290795** | 0.5034782 |
| IL-10 | **0.014964202** | 0.16080835 | 0.5034782 |
| IFN-g | **0.003330498** | 0.16080835 | 0.5034782 |
| TNF-a | 0.342512968 | 0.64712721 | 0.5034782 |
| GM-CSF | **0.014964202** | 0.23365079 | 0.5549938 |
| MIP-1a | **0.014964202** | 0.33591546 | 0.6287417 |
| P-Selectin | 0.911797181 | 0.31187418 | 0.6506653 |
| IL-1b | **0.014964202** | 0.44390222 | 0.6506653 |
| IL-6 | **0.004735736** | **0.03582795** | 0.6506653 |
| MCP-1 | **0.017686529** | **0.03582795** | 0.7017702 |
| IL-12p70 | **0.004735736** | 0.13858759 | 0.7275608 |
| IP-10 | **0.020985207** | **0.02013466** | 0.7765527 |
| IL-13 | 0.176113576 | 0.44390222 | 0.8652679 |
| IL-4 | **0.002298116** | 0.0563317 | 0.9074153 |
| IL-8 | **0.002298116** | **0.00938647** | 0.9082021 |
| MIP-1b | 0.176113576 | 0.33591546 | 0.9228697 |
| IL-1a | **0.038407413** | 0.07118931 | 0.9228697 |
