## Supplementary Table 2 for "Host-Microbial Interactions in Systemic Lupus Erythematosus and Periodontitis"

| Bacterial species | Non-periodontitis | | | Periodontitis | | |
|---|---|---|---|---|---|---|
| | SLE-I vs Control | SLE-A vs Control | SLE-A vs SLE-I | SLE-I vs Control | SLE-A vs Control | SLE-A vs SLE-I |
| C. ochracea | 0.848339 | **0.01492794** | **0.02985588** | 0.74233937 | **0.002085704** | **0.021098226** |
| S. intermedia | 0.6890417 | 0.43581042 | **0.04651062** | 0.21095715 | 0.398088605 | 0.656457659 |
| A. odontolyticus | 0.6890417 | 0.24663775 | **0.04651062** | 0.90136976 | 0.636131812 | 0.501481566 |
| F. polymorphum | 0.848339 | 0.24663775 | **0.04651062** | 0.90136976 | 0.235219839 | 0.107299566 |
| N. mucosa | 0.6890417 | 0.10404839 | 0.05495359 | 0.33780588 | 0.526837352 | 0.122425841 |
| T. socranskii | 0.848339 | 0.21356998 | 0.05495359 | 0.14262184 | 0.971063279 | 0.189737104 |
| E. saburreum | 0.6890417 | 0.43581042 | 0.05495359 | 0.90136976 | 0.559984792 | 0.467374925 |
| F. nucleatum | 0.9121669 | **0.04610472** | 0.05495359 | 0.13744175 | **0.008497623** | 0.259268922 |
| S. anginosus | 0.890477 | 0.09317978 | 0.06091701 | 0.42258803 | 0.125491208 | 0.533258614 |
| A. israelli | 0.848339 | 0.24663775 | 0.06091701 | 0.82244714 | 0.080678122 | 0.118102598 |
| V. parvula | 0.848339 | 0.22927351 | 0.07298449 | 0.74233937 | 0.559984792 | 0.259268922 |
| P. nigrescens | 1 | 0.43743455 | 0.0814709 | 0.90136976 | **0.008497623** | **0.040811022** |
| A. naeslundii | 0.6890417 | 0.24663775 | 0.08645407 | 0.09403132 | 0.411939204 | 0.070552457 |
| S. sanguinis | 0.7778798 | **0.01492794** | 0.08645407 | 0.95123914 | **0.049126921** | 0.052398425 |
| A. gerencseriae | 0.7778798 | **0.01686883** | 0.08645407 | **0.03353232** | 0.225474714 | **0.004816609** |
| P. micra | 0.6890417 | 1 | 0.12448247 | 0.13744175 | 0.360561317 | 0.423826251 |
| S. oralis | 0.7778798 | 0.34461457 | 0.16632558 | **0.01964188** | 0.235219839 | **0.011760555** |
| T. denticola | 0.848339 | **0.04610472** | 0.16856461 | 0.69093326 | 0.243334593 | 0.118102598 |
| C. gingivalis | 0.9823701 | 0.24663775 | 0.16856461 | 0.13744175 | **0.008497623** | 0.322213951 |
| E. nodatum | 0.6890417 | 0.92541297 | 0.1786198 | 0.1663759 | 0.559984792 | 0.423826251 |
| S. noxia | 0.848339 | 0.24227481 | 0.1786198 | **0.04182756** | 0.676795945 | 0.063845606 |
| C. showae | 0.7778798 | 0.43743455 | 0.19546746 | 0.74233937 | 0.411939204 | 0.656812785 |
| P. acnes | 0.7778798 | 1 | 0.21232112 | 0.14262184 | 0.554029415 | 0.289936529 |
| A. actinomycetemcomitans | 0.9823701 | 0.43743455 | 0.27953117 | 0.75449744 | 0.987603002 | 0.446864473 |
| A. viscosus | 0.9793132 | 0.15779199 | 0.29902871 | 0.69133769 | 0.83228158 | 0.322213951 |
| S. constellatus | 0.7778798 | 1 | 0.39097177 | 0.14262184 | 0.559984792 | 0.322213951 |
| P. melaninogenica | 0.7778798 | 1 | 0.39097177 | 0.94997833 | 0.962745528 | 0.750274161 |
| C. gracilis | 0.7778798 | 0.6960154 | 0.40467312 | 0.19575829 | 0.962745528 | 0.070552457 |
| T. forsythia | 0.9823701 | 0.24663775 | 0.44836274 | 0.46158523 | **0.008497623** | **0.040811022** |
| F. vincentii | 0.6890417 | 0.43743455 | 0.44836274 | 0.13744175 | 0.158864459 | 0.583094407 |
| C. rectus | 0.848339 | 1 | 0.51577983 | 0.33780588 | 0.875148401 | 0.322213951 |
| S. gordonii | 0.9823701 | 0.40569571 | 0.51577983 | 0.14262184 | **0.009475285** | 0.515704124 |
| S. mitis | 0.9823701 | 0.44502385 | 0.5922647 | 0.67763882 | 0.254528182 | 0.871096819 |
| F. periodonticum | 0.6890417 | 0.15779199 | 0.61351131 | 0.13744175 | 0.093974649 | 0.572420173 |
| C. sputigena | 0.7778798 | 0.43581042 | 0.85049649 | 0.14262184 | 0.360561317 | 0.259268922 |
| G. morbillorum | 0.7778798 | 0.26749601 | 0.85703994 | 0.14262184 | 0.411939204 | 0.322213951 |
| P. intermedia | 0.8680691 | 0.43743455 | 0.87412417 | 0.94997833 | 0.656603895 | 0.750274161 |
| P. gingivalis | 0.848339 | 0.44648528 | 0.89520858 | 0.13744175 | 0.398088605 | 0.322213951 |
| L. bucallis | 0.848339 | 0.43743455 | 0.99546146 | 0.1663759 | 0.342512994 | 0.509484631 |
| E. corrodens | 0.848339 | 0.44648528 | 1 | 0.36337257 | 0.962745528 | 0.190158017 |
