## Supplementary Table 3 for "Host-Microbial Interactions in Systemic Lupus Erythematosus and Periodontitis"

| Bacterial species | Non-periodontitis | | | Periodontitis | | |
|---|---|---|---|---|---|---|
| | SLE-I vs Control | SLE-A vs Control | SLE-A vs SLE-I | SLE-I vs Control | SLE-A vs Control | SLE-A vs SLE-I |
| C. ochracea | 1 | 0.9146953 | 1 | 1 | **0.002085704** | 1 |
| S. intermedia | 1 | 1 | 0.8693035 | 0.4982064 | 0.398088605 | 0.7962139 |
| A. odontolyticus | 1 | 0.9146953 | 0.8693035 | 1 | 0.636131812 | 1 |
| F. polymorphum | 1 | 1 | 0.8693035 | 1 | 0.235219839 | 1 |
| N. mucosa | 1 | 0.8187469 | 1 | 1 | 0.526837352 | 1 |
| T. socranskii | 1 | 1 | 1 | 0.9813349 | 0.971063279 | 1 |
| E. saburreum | 1 | 1 | 1 | 1 | 0.559984792 | 1 |
| F. nucleatum | 1 | 1 | 1 | 1 | **0.008497623** | 0.7962139 |
| S. anginosus | 1 | 0.8187469 | 1 | 0.9813349 | 0.125491208 | 1 |
| A. israelli | 1 | 1 | 1 | 1 | 0.080678122 | 1 |
| V. parvula | 1 | 1 | 1 | 1 | 0.559984792 | 1 |
| P. nigrescens | 1 | 1 | 1 | 1 | **0.008497623** | 1 |
| A. naeslundii | 1 | 1 | 1 | 1 | 0.411939204 | 1 |
| S. sanguinis | 1 | 0.8187469 | 1 | 1 | **0.049126921** | 1 |
| A. gerencseriae | 1 | 1 | 1 | 1 | 0.225474714 | 1 |
| P. micra | 1 | 1 | 1 | 1 | 0.360561317 | 1 |
| S. oralis | 1 | 1 | 1 | 0.8203875 | 0.235219839 | 0.7252702 |
| T. denticola | 1 | 0.8187469 | 1 | 1 | 0.243334593 | 1 |
| C. gingivalis | 1 | 1 | 1 | 0.9813349 | **0.008497623** | 1 |
| E. nodatum | 1 | 1 | 1 | 0.4982064 | 0.559984792 | 1 |
| S. noxia | 1 | 0.8187469 | 1 | 1 | 0.676795945 | 1 |
| C. showae | 1 | 0.9146953 | 1 | 1 | 0.411939204 | 1 |
| P. acnes | 1 | 1 | 0.8693035 | 0.4031895 | 0.554029415 | 0.7252702 |
| A. actinomycetemcomitans | 1 | 1 | 1 | 1 | 0.987603002 | 1 |
| A. viscosus | 1 | 0.8187469 | 1 | 1 | 0.83228158 | 1 |
| S. constellatus | 1 | 1 | 1 | 1 | 0.559984792 | 1 |
| P. melaninogenica | 1 | 1 | 0.8948576 | 1 | 0.962745528 | 0.7252702 |
| C. gracilis | 1 | 1 | 1 | 0.4982064 | 0.962745528 | 0.7962139 |
| T. forsythia | 1 | 1 | 1 | 1 | **0.008497623** | 1 |
| F. vincentii | 1 | 0.8187469 | 1 | 1 | 0.158864459 | 0.7252702 |
| C. rectus | 1 | 1 | 0.8693035 | 1 | 0.875148401 | 1 |
| S. gordonii | 1 | 0.9146953 | 1 | 0.4982064 | **0.009475285** | 1 |
| S. mitis | 1 | 0.8187469 | 0.8693035 | 1 | 0.254528182 | 0.7909436 |
| F. periodonticum | 1 | 0.8187469 | 1 | 0.4982064 | 0.093974649 | 0.7962139 |
| C. sputigena | 1 | 0.8187469 | 0.8693035 | 0.9813349 | 0.360561317 | 0.8428107 |
| G. morbillorum | 1 | 0.8187469 | 1 | 1 | 0.411939204 | 1 |
| P. intermedia | 1 | 0.9146953 | 1 | 1 | 0.656603895 | 1 |
| P. gingivalis | 1 | 0.8187469 | 1 | 0.4982064 | 0.398088605 | 1 |
| L. bucallis | 1 | 0.9146953 | 1 | 0.4982064 | 0.342512994 | 0.7962139 |
| E. corrodens | 1 | 0.8187469 | 1 | 0.4982064 | 0.962745528 | 1 |
